# COREMIC: a web-tool to search for a root-zone associated CORE MICrobiome

**DOI:** 10.1101/147009

**Authors:** Richard R. Rodrigues, Nyle C. Rodgers, Xiaowei Wu, Mark A. Williams

## Abstract

Microbial diversity on earth is extraordinary, and soils alone harbor thousands of species per gram of soil. Understanding how this diversity is sorted and selected into habitat niches is a major focus of ecology and biotechnology, but remains only vaguely understood. A systems-biology approach was used to mine information from databases to show how it can be used to answer questions related to the core microbiome of habitat-microbe relationships. By making use of the burgeoning growth of information from databases, our tool “COREMIC” meets a great need in the search for understanding niche partitioning and habitat-function relationships. The work is unique, furthermore, because it provides a user-friendly statistically robust web-tool (http://coremic2.appspot.com), developed using Google App Engine, to help in the process of database mining to identify the “core microbiome” associated with a given habitat. A case study is presented using data from 31 switchgrass rhizosphere community habitats across a diverse set of soil and sampling environments. The methodology utilizes an outgroup of 28 non-switchgrass (other grasses and forbs) to identify a core switchgrass microbiome. Even across a diverse set of soils (5 environments), and conservative statistical criteria (presence in more than 90% samples and FDR *q*-val < 0.05% for Fisher’s exact test) a core set of bacteria associated with switchgrass was observed. These included, among others, closely related taxa from *Lysobacter spp., Mesorhizobium spp*, and *Chitinophagaceae*. These bacteria have been shown to have functions related to the production of bacterial and fungal antibiotics and plant growth promotion. COREMIC can be used as a hypothesis generating or confirmatory tool that shows great potential for identifying taxa that may be important to the functioning of a habitat (e.g. host plant). The case study, in conclusion, shows that COREMIC can identify key habitat-specific microbes across diverse samples, using currently available databases and a unique freely available software.

## 1. Introduction

Microbial diversity on earth is extraordinary, and soils alone harbor thousands of species per gram (Hughes et al., 2001). Understanding how this diversity is sorted and selected into habitat niches is a major focus of ecology and biotechnology, but remains only vaguely understood. The advent of next-generation sequencing technologies now allow for the potential to make great leaps in the study of microbe-habitat relationships of highly diverse microbial communities and environments. The identity and functions of this overwhelming multitude of microbes are in the beginning stages of being described, and are already providing insights into microbial impacts on plant and animal health (Berg, 2009; Evans and Schwarz, 2011; Clemente et al., 2012). Making use of the overwhelming amount of information on microbial taxa and habitats has enormous potential for use to further understand microbial-habitat relationships. Thus, the advent of new methods and approaches to utilize this data and describe microbiomes will benefit microbial ecology and biotechnology.

Though variations exist, a core microbiome can be defined, conceptually, using Venn diagrams, where over-lapping circles and non-overlapping areas of circles represent shared and non-shared members of a habitat, respectively (Shade and Handelsman, 2012). Typically, microbiomes identified in this manner are not statistically evaluated, or by nature, seek to answer specific hypothesis that are specific to an experiment. For example, studies often identify microbes associated with different plant growth stages, species, cultivars, and locations but rarely, if at all, mine databases or perform meta-analysis to statistically identify microbiomes across studies and experimental conditions (Chaudhary et al., 2012; Liang et al., 2012; Mao et al., 2013; Mao et al., 2014; Hargreaves et al., 2015; Rodrigues et al., 2015; Jesus et al., 2016; Rodrigues et al., 2017). Describing differences due to treatment or habitat conditions are informative in their own right, however, extending this framework to include an easy to use, and statistically robust tool to help in the mining of data from underutilized and burgeoning databases (e.g. the National Center for Biotechnology Information (NCBI), Riboso-mal Database Project) can help transform the ecological study of microbes in their natural environment. Using the vast and growing databases of organism and habitat metadata will allow for both the testing and development of hypotheses associated with habitat-microbe relationships that were not formerly possible.

To address the challenges described above, we developed COREMIC - a novel, easy to use, and freely available web tool to identify the “core microbiome”, of any well-defined habitat (e.g. plant root-zone) or niche (Shade and Handelsman, 2012). This straightforward approach is a novel and powerful way to complement existing analysis (e.g. indicator species analysis (ISA) (Dufrene and Legendre, 1997)) by allowing for the use of data that is now overflowing among freely available databases. It seeks to determine the core set of microbes (core microbiome) that are explicitly associated with a host system or habitat. The ability to identify core microbiomes at this scale has great potential to describe host-microbe interactions and habitat preferences of microbes.

A meta-analysis based case study was performed, combining diverse sequencing datasets derived from NCBI, to test for the occurrence of a core microbiome in the rhizosphere (root-zone) of switchgrass. Switchgrass is a US-native, perennial grass studied by many researchers, and thus has a growing database to mine for genetic information. Its widespread study is likely a result of its bioenergy potential, and the capacity of the grass to grow on marginal lands not dedicated to crops. Studies have identified different bacteria found in the root-zones of switchgrass (Jesus et al., 2010; Mao et al., 2011; Chaudhary et al., 2012; Liang et al., 2012; Mao et al., 2013; Bahulikar et al., 2014; Mao et al., 2014; Werling et al., 2014; Hargreaves et al., 2015; Jesus et al., 2016; Rodrigues et al., 2017), however, there has been no integrative study of different datasets identifying the core microbiome in switchgrass rhizospheres. It is thus proposed to identify host-habitat relationships as a proof of concept for a core microbiome. In this paper we utilize a plant host to define a habitat, but theoretically any habitat and associated organisms could make use of COREMIC and its approach to identify a core microbiome.

## 2. Material and methods

### 2.1. Datasets used in the study

A diverse set of data composed of 61 samples from two different published datasets and collected from multiple locations (Jesus et al., 2016; Rodrigues et al., 2017) were used for this study. Data were obtained from the NCBI and selected based on the availability of the raw (16S rRNA) sequence data of root-zone bacteria from switchgrass and that for an outgroup of reference (native and/or other grasses) plants.

The dataset “Jesus 2016”(Jesus et al., 2016), PRJEB6704, compared the rhizosphere soil microbial communities associated with restored prairie with three grass crops, namely corn, switchgrass, and mixed prairie grasses. The grasses were grown in fields of Michigan and Wisconsin and were harvested after two and ten years. The V6-V8 region of the 16S rRNA gene was amplified and sequenced using the Roche 454 pyrosequencing. In our study, we used a total of 43 samples (3 each from corn, switchgrass, mixed grasses (2 yrs. only), and restored prairie grasses grown in Wisconsin and Michigan, and sampled after 2 and 10 years. Switchgrass grown in Michigan, composed of 4 samples, were collected following 10 years of plant growth.

The dataset “Rodrigues 2017”(Rodrigues et al., 2017), PRJNA320123, compared the root-zone soil microbial communities associated with switchgrass cultivars: “Alamo” and “Dacotah”. The switchgrass were grown in the greenhouse using soil derived from plots growing Switchgrass (>7 years) near Blacksburg, VA. Switchgrass rhizosphere bacteria were sampled at three different growth stages. The V3-V4 region of the 16S rRNA gene was amplified and sequenced using Illumina MiSeq sequencing. In our study, we used a total of 18 switchgrass samples for Alamo (A) and Dacotah (D) from stages V2 and E3 (4 AV2, 4 DV2, 5 AE3, 5 DE3 = 18).

Overall, these datasets served as a diverse resource (relevant differences are summarized in Figure 1) to compare the root-zone bacteria and identify core-bacteria associated with switchgrass.

**Figure 1:**
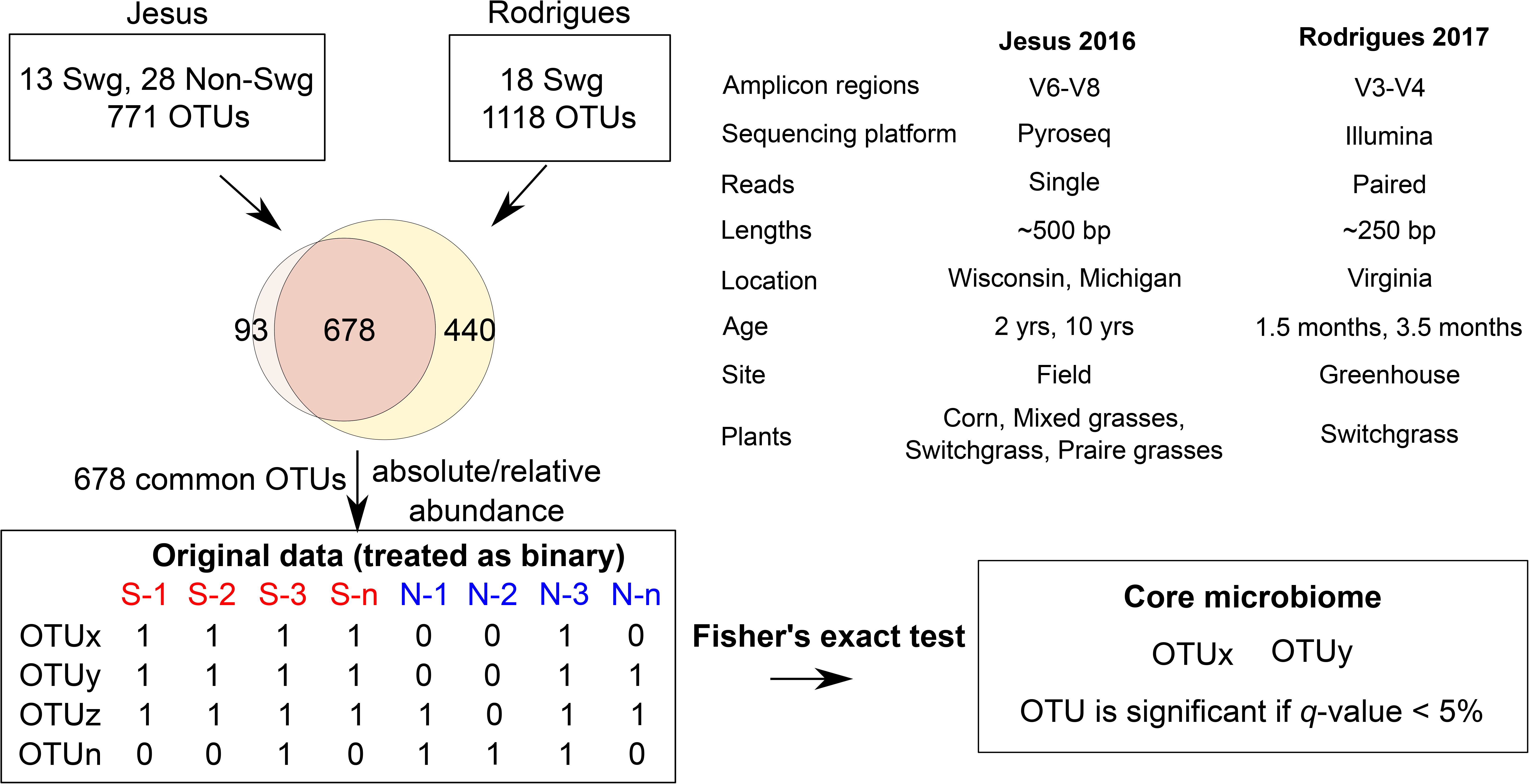
The COREMIC approach. The workflow indicating the Jesus 2016 and Rodrigues 2017 datasets and differences between them, and the methodology used to identify core microbiome. Switchgrass and other grasses are indicated by “Swg” and “Non-Swg,” respectively.

### 2.2. Sequence data analysis and picking of Operational Taxonomic Units (OTU)

For the Rodrigues 2017 dataset, the OTU table was obtained from previously performed analysis (Rodrigues et al., 2017). For the Jesus 2016 dataset, quality score (25) and read lengths (150) thresholds were enforced using cutadapt (1.8.1) (Martin, 2011) and an open reference OTU picking (enable_rev_strand_match True) was performed in QIIME v1.8.0 (Caporaso et al., 2010), as previously described (Rodrigues et al., 2015; Rodrigues et al., 2017), to allow comparison with the other dataset. Briefly, uclust (Edgar, 2010) was used to cluster reads into OTUs (97% sequence similarity) and assign taxonomy against the Greengenes reference database version 13.8 (DeSantis et al., 2006; McDonald et al., 2012). Two samples from the Jesus 2016 dataset were removed from downstream analysis due to very few sequences assigned to OTUs.

### 2.3. Combining two datasets

Within each OTU table, sequences assigned to identical OTUs in a sample were summed to retain unique taxa. The common (678) OTUs from the two datasets were selected, converted to biom format and used for further analyses (Figure 1). The data table was filtered and rarefied using a sequence threshold of 1150, and the beta diversity was calculated using Bray-Curtis (Beals, 1984) distance and visualized using Principal Coordinate Analysis (Gower, 2005). Multivariate data analysis methods of MRPP (Mielke, 1984), Permanova (Anderson, 2001) and ANOSIM (Clarke, 1993) were used to identify whether the plant type (switchgrass versus non-switchgrass) were associated with different bacterial communities.

### 2.4. Core microbiome analysis

To find the set of core OTUs, the samples in the combined OTU table (original data) were first divided into the interest group samples (switchgrass) and out-group samples. The abundance values for each OTU in each sample are then converted to binary (present/absent) values based on whether they are zero or nonzero. For each OTU a one-tailed Fisher’s Exact Test was used to calculate a *p*-value testing whether an OTU was present in a significantly higher portion in the interest in-group (Switchgrass) compared to the out-group samples (numerous other grass species).

These *p*-values were corrected for multiple-testing using Benjamini Hochberg. The OTUs with a *q*-value < 0.05 were then selected to only the OTUs that are present in at least 90% of the interest group samples. Uninformative OTUs (e.g., k_Bacteria;p_;c_;o_;f_;g_;s_) were filtered out and the remaining OTUs were candidates for the core microbiome.

### 2.5. Implementation of COREMIC

COREMIC and the datasets are available at http://coremic2.appspot.com. Its code is available on github (https://github.com/richrr/coremicro). The web-tool was developed in Python 2.7, and is hosted on Google App Engine. Other requirements include GoogleAppEnginePipeline 1.9.22.1, pyqi 0.3.1, requests 2.10.0, requests-toolbelt 0.6.2, mailjet-rest 1.2.2, biom-format 1.1.2, ete3 3.0.0 (for tree generation—see below for details), webapp2 2.5.2, numpy 1.6.1, matplotlib 1.2.0, jinja2 2.6, ssl 2.7. COREMIC is accessible via any internet connected browser and emails the results to the user. The processing times with the default settings after uploading the data are provided in Table S1.

A custom python script generates a phylogenetic tree using the taxonomic labels for each OTU displaying the relationship between the core OTUs obtained from the group of interest and the out-group. This tree is generated using the ete3 3.0. 0 library.

## 3. Results

After quality filtering, a total of 319,821 reads were obtained from the Jesus 2016 dataset (mean 461.45 and std. dev. 69.34). Two samples with very few (48 and 75) counts were removed; each of the remaining samples had more than 1150 sequences assigned to OTUs. The number of OTUs in the Jesus 2016 and Rodrigues 2017 datasets was 771 and 1118, respectively. The combined dataset had 678 OTUs, 31 switchgrass and 28 non-switchgrass (other grasses) samples.

The bacterial communities in switchgrass and grasses from the combined dataset were significantly different (Permanova, MRPP, and ANOSIM *p*-values < 0.01) and as can be observed using the PCoA plot using the Bray-Curtis dissimilarity metric (Figure 2). These differences were apparent despite significant difference across datasets (Permanova, MRPP, and ANOSIM *p*-values < 0.01); which could be the result, for example, of the heterogeneity of the data set related to climate, soil type-condition, growth conditions, and plant age. In this regard, at the phylum level, Mann Whitney test identified Bacteroidetes and Verrucomicrobia had significantly greater (*p*-value < 0.05) relative abundance in switchgrass, whereas, Gemmatimonadetes were more abundant in other grasses (Figure S1).

**Figure 2:**
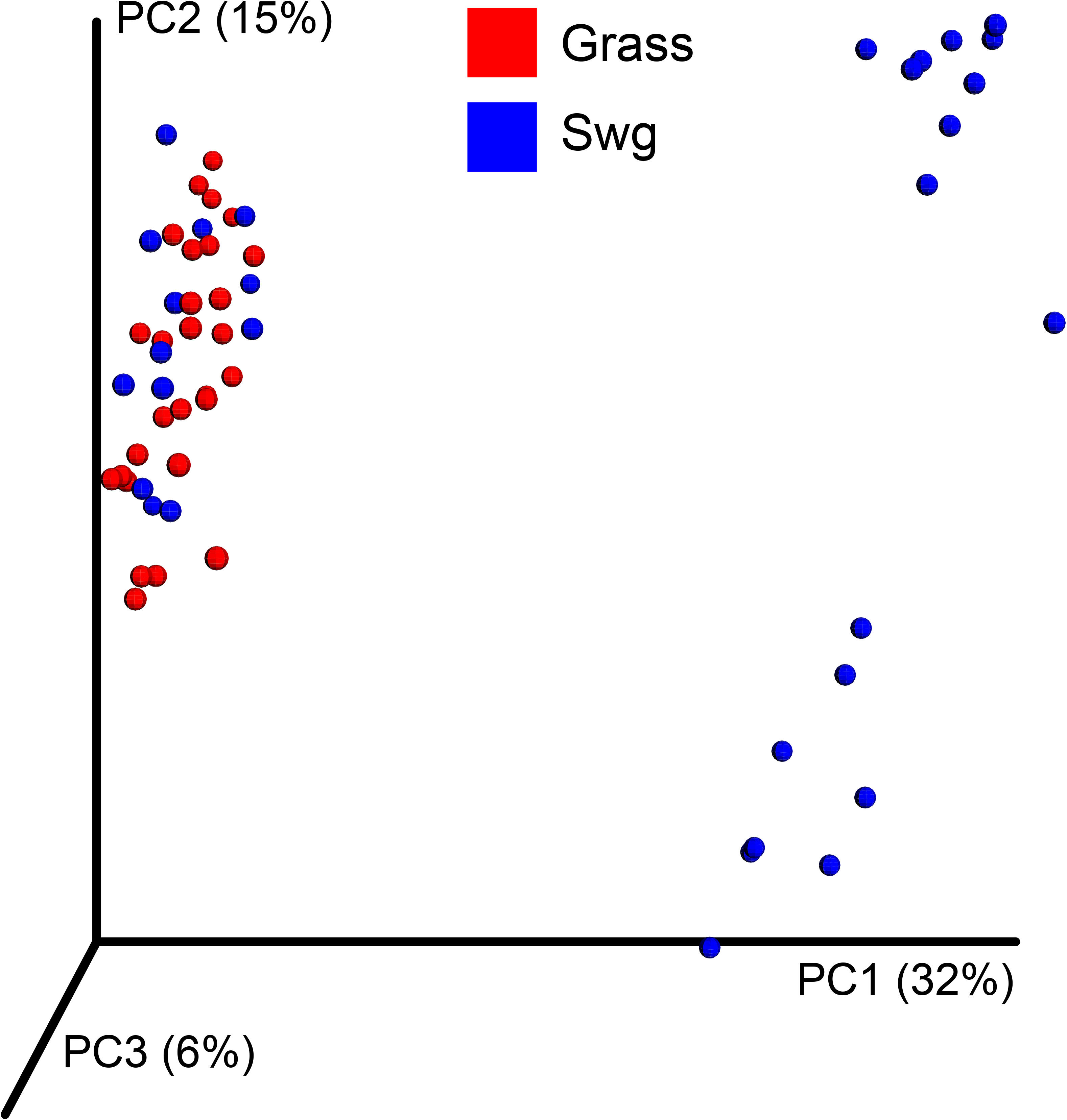
Beta-diversity of the combined dataset. PCoA plot showing Bray-Curtis dissimilarities for bacterial communities at the OTU level in switchgrass (blue colored) and other grasses (red colored).

We used a very conservative criterion of >90% threshold i.e., an OTU has to be present in at least 90% of switchgrass samples and observed five OTUs with FDR *q*-values < 0.05 (Table 1). The relative abundance and a phylogenetic tree exhibiting their relationship with the core-OTUs from the non-switchgrass samples is shown in Figure S2 and Figure S3, respectively. Despite the enormous variability across the many different sampling locations, there is support for the occurrence of a core microbiome in the root-zone of switchgrass.

**Table 1:** Bacterial OTUs associated with switchgrass.

| OTU | present(%) |
| --- | --- |
| p_Proteobacteria;c_Gammaproteobacteria;o_Xanthomonadales;f_Xanthomonadaceae;g_Lysobacter;s_ | 100 |
| p_Planctomycetes;c_Planctomycetia;o_B97;f_;g_;s_ | 96.8 |
| p_Bacteroidetes;c_[Saprospirae];o_[Saprospirales];f_Chitinophagaceae | 96.8 |
| p_Proteobacteria;c_Alphaproteobacteria;o_Rhizobiales;f_Phyllobacteriaceae;g_Mesorhizobium;s_ | 90.3 |
| p_Proteobacteria;c_Gammaproteobacteria;o_Legionellales;f_;g_;s_ | 90.3 |

The core bacterial OTUs those were significantly ( $q$ -value $< 0.05$ ) associated with switchgrass, calculated using presence/absence data and present in $>90\%$ switchgrass samples.

## 4. Discussion

The case study showed how COREMIC can identify key habitat-specific microbes across diverse samples, using currently available databases and a unique freely available software. The core set of bacteria associated with switchgrass included, among others, closely related taxa from *Lysobacter spp., Mesorhizobium spp*, and *Chitinophagaceae*. The functional relevance of these bacteria related to switchgrass is unknown, but it is notable that these bacteria have been shown to produce bacterial and fungal antibiotics and promote the growth of plants (Kaneko et al., 2000; Kilic-Ekici and Yuen, 2004; Weir et al., 2004; Islam et al., 2005; Jochum et al., 2006; Ji et al., 2008; Park et al., 2008; Nandasena et al., 2009; Yin, 2010; Bailey et al., 2013; Degefu et al., 2013; Guerrouj et al., 2013; Madhaiyan et al., 2015). The analyses from the highly diverse data sets thus provided information that helps to greatly narrow down possibilities and thus set the stage for testing, using controlled studies, how the core microbiota potentially support or antagonize the function of a native grass. This novel toolkit is simple to use and supports use by a broad range of biological scientists, and is particularly relevant to those with expertise in their field but with limited bioinformatics background. Overall, in a dataset derived from a complex and diverse set of habitats and ecosystems, this tool was shown to pinpoint microbiota of the microbiome that might have important functional implications within their habitat or host.

### 4.1. Methodological considerations in the use of COREMIC

COREMIC performs a complementary analysis different from that of existing methods by using presence/absence data. For two groups (A and B) it checks whether (pre-determined percentage of) samples from group A have a non-zero value for the OTU. This allows scientists to operate without making assumptions about the PCR-based OTU relative abundances. This is considered a potential advantage of the method because it is unknown whether relative abundance of sequence data is representative of true relative differences between communities. Further research, in this regard, will be aimed towards investigating other measures of OTU “presence”, namely the extent of exclusivity, consistency, or abundance of the group that is eventually determined to be a core microbiome.

Sampling plots used in this study were located across a range of diverse environments to help create a backdrop of heterogeneity. While this diversity of habitat conditions ignores the potential for microbe-environment interactions that might be important for the plant-microbial relationship, it has the advantage of being a conservative approach with high veracity for defining a core microbiome regardless of habitat heterogeneity. The locations from which samples were grown (Michigan, Wisconsin, Virginia) were treated as independent to help isolate the overall habitat effect of switchgrass (Werling et al., 2014; Jesus et al., 2016). When the effects of habitat are thought to be habitat specific, researchers can take this into account during the design and analysis using COREMIC.

It is notable that the representation of an outgroup (multiple non-switchgrass species) is an important criteria and choice made by researchers, and is an approach that has both advantages and caveats. By definition, a habitat is defined by its differences from that of other habitats, and therefore the use of the outgroup is an important choice. A counterargument for the current dataset might argue for exclusion of breeding lines of a cultivated grass (maize) as being unrepresentative of the grass outgroup. In our case, it was thought, *a priori*, that a diverse set of grasses would provide the best comparison; and no compelling argument was found that supported the exclusion of maize from the analysis. An implicit assumption was also made that the taxonomy of plant species (root-zone habitats) play an important role in determining root-zone microbial communities, an approach supported by extensive findings that different grass species associate with different microbial communities (Kuske et al., 2002; Kennedy et al., 2004; Berendsen et al., 2012; Chaudhary et al., 2012; Turner et al., 2013). So although there is a need for careful consideration of the experimental questions of interest when using COREMIC, this is a common, if not ubiquitous foundation of all experimentation and hypothesis testing. The results provide a statistically valid approach using freely available software to describe and define a core microbiome of switchgrass.

The choice of the outgroup, furthermore, for determining a core microbiome is amenable to choice using deductive reasoning but ultimately limited by available data. This issue almost certainly limits inclusion of many functionally important rhizosphere microbes that could affect the growth of switchgrass. In this study, the proof of concept utilized a conservative approach to highlight the methodology across a diversity of geographies, soil types, and plant ages. The COREMIC tool as well as the multiple methods for defining a core microbiome (e.g., QIIME (Caporaso et al., 2010),

ISA (Dufrene and Legendre, 1997)) will always be defined by the expertise, and the nature of the hypotheses defined and defended by individual researchers.

### 4.2. Core Microbes

The individual datasets described in this study had previously focused on identifying abundant microbes and differences due to experimental conditions. The current meta-analysis goes a step further to find common microbiota that are associated with switchgrass across the diverse experimental conditions. The members of the *Lysobacter* genus, an identified core microbe of switchgrass, are known to live in soil and have been shown to be ecologically important due to their ability to produce exo-enzymes and antibiotics (Reichenbach, 2006). Their antimicrobial activities against bacteria, fungi, unicellular algae, and nematodes have been described (Islam et al., 2005; Jochum et al., 2006; Park et al., 2008; Yin, 2010). Strains of this genus, for example, have been used for control of diseases caused by bacteria in rice (Ji et al., 2008) and tall fescue (Kilic-Ekici and Yuen, 2004). Reports of their function thus support the idea that they may play an important role in switchgrass growth and survival. The core microbiome results thus support further research into the role played by this bacterium in the switchgrass rhizosphere.

Similarly, members of the *Mesorhizobium* genus are well-known diazotrophs (Kaneko et al., 2000) and previously shown to be symbiotically associated with switchgrass (DeAngelis et al., 2010; Bahulikar et al., 2014) and legumes (Weir et al., 2004; Nandasena et al., 2009; Degefu et al., 2013; Guerrouj et al., 2013). Another identified core microbiome taxa, soil-dwelling members of the *Chitinophagaceae* family are known to have β-glucosidase (Bailey et al., 2013) and Aminocyclopropane-1-carboxylate (ACC) deaminase activities and ability to produce indole-3-acetic acid (IAA) (Madhaiyan et al., 2015). These molecules and enzymes are well known for their effects on plant growth (Zhao, 2010; Van de Poel and Van Der Straeten, 2014). The capacity to degrade cellulose might provide additional and readily available options to aid survival of these bacteria near switchgrass root zones during times of environmental stress. ACC deaminase and IAA production, in contrast, are potent plant growth modulators (Glick, 2014) that could play a role in plant productivity and survival, especially under conditions of plant physiological stress. Though these examples above would need further study, they provide consistent examples describing how a core microorganism could play a role in determining plant function and growth. The power of the approach stems from the ability to identify the core microbes associated with a plant (or other habitat), and that can, with veracity, narrow down potentially important core microbes from otherwise hyperdiverse samples.

From a technological standpoint, it is important to put the current approach into context with research before the metagenomics era. The search and identification of antagonistic plant growth promoting microbes has previously been tedious and labor intensive. Screenings of hundreds of microbes were used to cultivate and identify candidate microbes that might support (or deter) plant growth. In the case of beneficial microbes, even when identified under greenhouse conditions, the beneficial effects rarely translated into plant supportive growth under field growth conditions (Babalola, 2010; Hayat et al., 2010). With the aid of hindsight and new knowledge suggesting the importance of the soil habitat and root-soil interactions in the development of growth promoting plant-microbial relationships, the approach used in this study reverses the focus (from top-down to bottom-up) to search for microbes that appear to already be naturally well-adapted to the root-soil habitats of interest (Trabelsi and Mhamdi, 2013; Souza et al., 2015). This process streamlines the search for suitable microbes from a daunting pool of thousands of bacterial taxa. Bacteria and fungi with well-known partnerships with members of the core microbiome, it would be expected, to be more readily adaptable to their native environment. Indeed, the concept of adaptability to an environment has been shown to be true for many types of microbes across the environmental spectrum, and has given rise to the concept of the niche (Lennon et al., 2012). The COREMIC tool provides an alternative and logical approach to help mine available datasets, in the search for core microbiomes associated with habitats that are ecologically and agriculturally important.

### 4.3. Conclusions

The COREMIC tool, by helping to mine multiple datasets fills a major gap in the search for the core microbiome associated with a host or habitat. It allows for the development of a working hypothesis in the search for microbes well suited for a habitat or host-microbe interaction. It can also be used to confirm laboratory studies that have identified target microbes that might be important symbionts or thought to be associated with a specific habitat. In the case of plants, but not limited to them, the COREMIC approach can identify microbial targets that might be useful for plant growth promotion. An example of this would be the identification of diazotrophic bacteria that aid the growth of bioenergy grasses and help to serve the development of sustainable agricultural systems. This combined with the ongoing efforts of plant breeding and genetic modification would help to catalyze microbe-driven crop yield improvement while practicing environmental stewardship through reduced fertilizer use. Here we show the applicability of COREMIC in rhizosphere-associated microbes, but the overall concepts are translational across disciplines with interests in host-microbe and microbe-habitat relationships. The applicability of COREMIC for the identification of core genes and microbes has excellent potential to help understand the roles of microorganisms in complex and diverse microbial communities.

## Declarations

### Ethics approval and consent to participate

Not applicable.

## Consent for publication

Not applicable.

## Availability of data and materials

The datasets and results supporting the conclusions of this article are included within the article and supplementary files. COREMIC and the datasets are available at http://coremic2.appspot.com. An archived version of its code is available on github (https://github.com/richrr/coremicro). COREMIC and its code are freely available under the GPL license.

## Authors’ contributions

Conceived and designed the experiments: RRR MAW. Implemented software tools: RRR NCR. Performed the experiments: RRR NCR. Analyzed the data: RRR NCR XW MAW. Wrote the paper: RRR NCR XW MAW. All authors read and approved the final manuscript.

## Acknowledgements

The authors thank Dr. Roderick Jensen and James Wrenn for their suggestions in improving the manuscript. We thank Dr. James McClure for his help in the web-tool development. We also acknowledge BIOM, Google App Engine and their developers. The authors acknowledge Virginia Polytechnic Institute and State University's Open Access Subvention Fund. We thank Virginia Tech’s Genetics, Bioinformatics, and Computational Biology, Department of Horticulture for providing personnel funding. Research was also partially funded by grants from USDA-NIFA (2011-03815).

---

**Figure S1: Taxonomic summary of the relative abundance of bacterial phyla in the combined dataset.** The taxa and the labels are arranged as per total relative abundance across all samples, with the most abundant phyla at the bottom and the least abundant phyla at the top of the y-axis. Mann Whitney test was used to identify phyla with significantly different (p value < 0.05) relative abundance.

**Figure S2: Abundance of core microbiome of switchgrass.** The bar plot compares the relative abundance of switchgrass (red colored) core OTUs (90% threshold and q-value < 0.05) and non-switchgrass (yellow colored) samples.

**Figure S3: Core microbiome of switchgrass.** Phylogenetic tree showing relationships between core OTUs (90% threshold and q-value < 0.05) identified from switchgrass (blue colored) and non-switchgrass samples.

**Table S1:**
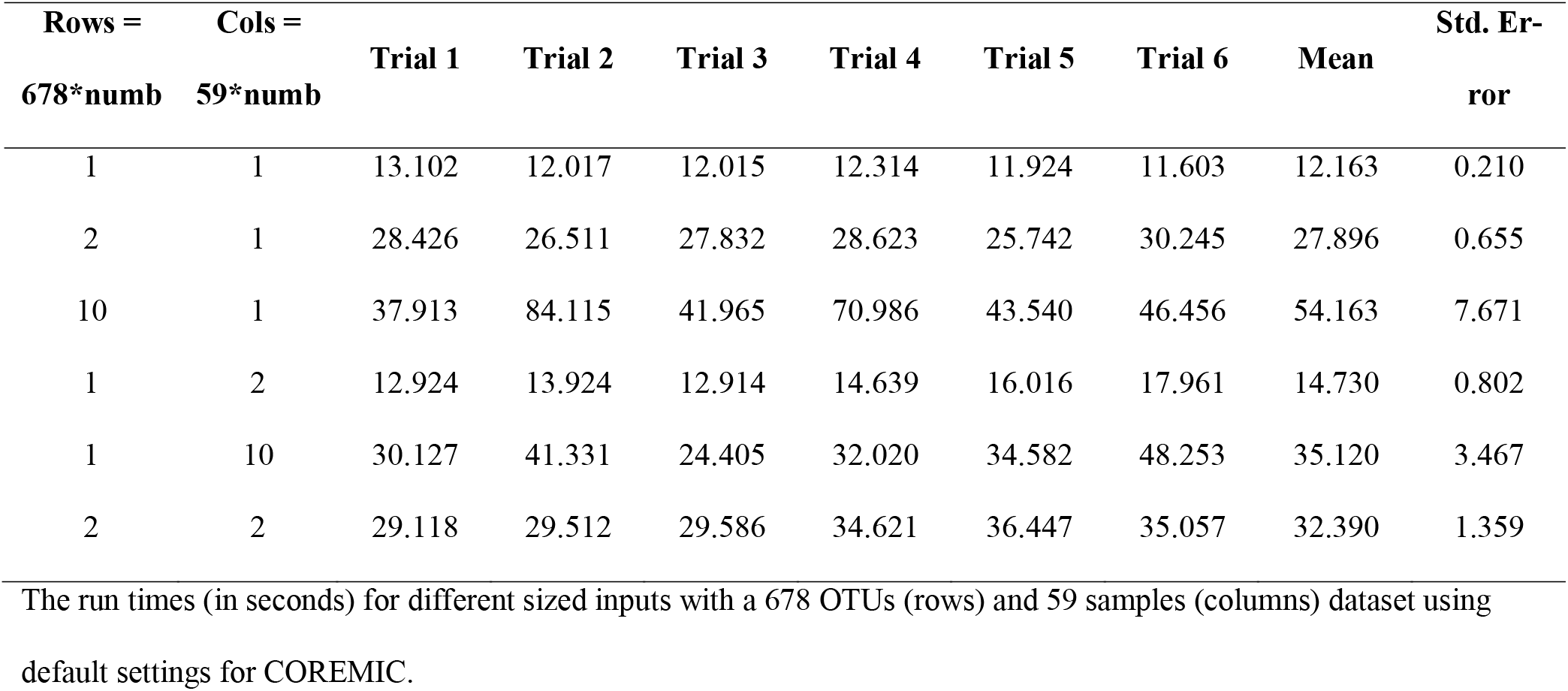
Processing times for COREMIC.

| Rows =<br>678*numb | Cols =<br>59*numb | Trial 1 | Trial 2 | Trial 3 | Trial 4 | Trial 5 | Trial 6 | Mean | Std. Error |
| --- | --- | --- | --- | --- | --- | --- | --- | --- | --- |
| 1 | 1 | 13.102 | 12.017 | 12.015 | 12.314 | 11.924 | 11.603 | 12.163 | 0.210 |
| 2 | 1 | 28.426 | 26.511 | 27.832 | 28.623 | 25.742 | 30.245 | 27.896 | 0.655 |
| 10 | 1 | 37.913 | 84.115 | 41.965 | 70.986 | 43.540 | 46.456 | 54.163 | 7.671 |
| 1 | 2 | 12.924 | 13.924 | 12.914 | 14.639 | 16.016 | 17.961 | 14.730 | 0.802 |
| 1 | 10 | 30.127 | 41.331 | 24.405 | 32.020 | 34.582 | 48.253 | 35.120 | 3.467 |
| 2 | 2 | 29.118 | 29.512 | 29.586 | 34.621 | 36.447 | 35.057 | 32.390 | 1.359 |

The run times (in seconds) for different sized inputs with a 678 OTUs (rows) and 59 samples (columns) dataset using default settings for COREMIC.

